# Feedback reliability modulates learning-related dynamics of midfrontal error monitoring

**DOI:** 10.64898/2026.10.30.762838

**Authors:** Magda Gawlowska

## Abstract

Learning from feedback requires the brain to evaluate both performance outcomes and the reliability of the information used for learning. This study examined the dynamics of neural performance monitoring during deterministic (100% reliable) and probabilistic (75% reliable) associative learning using two within-subject tasks with different stimulus materials. Behavioral performance improved under both feedback regimes, but learning was consistently more successful under deterministic feedback. Error-related negativity, feedback-related negativity, and phase-locked midfrontal theta became progressively more pronounced across deterministic learning, particularly in the image-image task, whereas corresponding changes were attenuated under probabilistic feedback. Induced theta showed the opposite pattern, decreasing across deterministic learning while remaining comparatively stable under probabilistic learning. These opposing theta trajectories suggest that phase-locked and induced activity reflect distinct components of learning-related neural adaptation that may be obscured when total theta power is considered alone. Overall, the findings indicate that feedback reliability shapes the progressive tuning of midfrontal performance-monitoring responses during learning.

## Introduction

Learning from feedback is fundamental to adaptive behavior, and a substantial body of research has identified the anterior cingulate cortex (ACC) as a central hub of the neural system that supports it. The ACC and adjacent medial frontal regions generate two well-characterized event-related potentials (ERPs) — the error-related negativity (ERN), peaking shortly after a self-detected response error, and the feedback-related negativity (FN), peaking after the delivery of negative feedback information, which index the system’s response to performance-relevant events [1]. At the level of oscillatory activity, these components are accompanied by transient increases in midfrontal theta-band (4–8 Hz) power, which has been characterized as a general substrate of cognitive control [2,3,4]. Recent theoretical accounts converge that this system actively integrates prediction-error signals with information about the structure and predictability of the environment to support adaptive behavioral adjustments [5,6].

A majority of the literature on this system has been built using probabilistic learning paradigms, in which the relationship between cues, responses, and outcomes is stochastic [7,8], while deterministic learning, where cue is reliably associated with a single correct response and feedback is always truthful, has been comparatively less examined. Prior work by the present author and colleagues established that in deterministic paired-associate learning, both the ERN and FN deepen progressively across learning stages, with corresponding increases in response- and feedback-related ACC activity [9,10]. These findings demonstrated that deterministic learning is accompanied by a clear neural trajectory of error-monitoring development, but they did not address the converse question: how does this trajectory differ when feedback is only partially reliable? Studies that have used feedback-validity manipulations have typically employed the deterministic-versus-probabilistic contrast as a secondary axis for probing individual differences such as aging [11,12,13,14], with learning dynamics often reduced to an early-versus-late comparison. To the author’s knowledge, a direct within-subject comparison of learning trajectories between fully reliable and partially misleading feedback environments, using a multi-measure characterization of error-monitoring dynamics across multiple stages, has not previously been undertaken.

Characterizing these dynamics in a theoretically informative way requires more than a single measure. The ERN and FN, although temporally localized, capture the phase-locked, time-domain component of midfrontal error-monitoring activity; richer information is available from the time–frequency domain, where total theta power and its phase-locked and induced sub-components index complementary aspects of the underlying neural response [8,15,16]. Recent work has further suggested that midfrontal theta and the FN are sensitive not only to feedback value, but also to its salience as a learning signal, including its motivational and attentional relevance for behavioral adjustment [17,18]. Together, these considerations motivate a multi-measure approach in which convergence across ERPs and theta-power measures can reveal how the error-monitoring system progressively tunes its responses to the reliability of a learning environment.

A further methodological consideration concerns the decomposition of midfrontal theta into phase-locked (evoked) and induced (non-phase-locked) sub-components. The phase-locked component reflects neural activity consistently time-locked to the eliciting event and underlies the ERPs, whereas the induced component captures oscillatory activity that varies in phase across trials and is typically associated with more sustained, non-stimulus-locked processing [19,16]. The two sub-components may therefore index dissociable neural processes as has been demonstrated in motor practice paradigms, in which induced frontal theta decreases across extensive training while evoked theta and the FN do not show the same decline [20]. Time–frequency decomposition along these lines has been applied in feedback-processing research using gambling and reward paradigms [21], but its application to learning trajectories under contrasting conditions of feedback reliability has not been thoroughly explored.

A final design consideration concerns the generality of any observed dynamics. The use of a single stimulus paradigm leaves open the possibility that observed effects may depend partly on the specific material being learned [22]. In the present study, two distinct paradigms were employed, which use markedly different stimulus material — abstract image–image pairs adapted from Gawlowska et al. [9] and face–name pairs adapted from Alm, Rolheiser and Olson [23]. These tasks differ in the cognitive characteristics of the material: the image–image paradigm (the PICTURE task) involves associative encoding with emotionally neutral material, whereas the face–name paradigm (the FACE task) additionally involves socially salient facial stimuli [24]. Performance-monitoring components such as the ERN and FN have been shown to be modulated by emotional and motivational context, with cognitive control itself increasingly understood as resting on an emotional foundation [25,26]. A consistent pattern of error-monitoring dynamics across both paradigms would therefore provide evidence for cross-paradigm generality and support the possibility of a domain-general mechanism operating across materially different associative-learning contexts.

The present study addresses these considerations through a within-subject EEG experiment in which participants completed both paradigms under deterministic (DET, 100% reliable feedback) and probabilistic (PRO, 75% reliable feedback) conditions, across multiple within-subject learning stages. Five complementary measures of error-monitoring activity — the ERN, FN, total midfrontal theta power, and its phase-locked and induced sub-components — were examined as a function of feedback reliability, response/feedback type, and learning stage. It was predicted that the dynamics of error-monitoring responses (ERN, FN, and phase-locked theta) would diverge between the deterministic and probabilistic conditions across learning stages, with progressive amplification in the deterministic condition on error and negative-feedback trials and a comparatively static profile in the probabilistic condition. A further prediction was that the overall pattern of reliability-dependent error-monitoring dynamics would be similar across both associative-learning paradigms, providing evidence for cross-paradigm consistency and supporting the possibility of a domain-general mechanism of error-driven learning. Moreover, the study explored whether the phase-locked and induced sub-components of midfrontal theta would show dissociable trajectories across learning, given recent evidence that they may index distinct neural processes [20,16].

## Results

### Behavioral performance

#### Learning dynamics: accuracy and reaction time

Learning was quantified using d-prime (d′) from signal detection theory [27], with the Hautus [28] log-linear correction, at each learning stage and complemented by mean reaction times (RTs) at the same stages. Two-way repeated-measures ANOVAs (Condition × Learning Stage) were applied to both measures, separately for the two tasks (Fig. 1).

**Figure 1.**
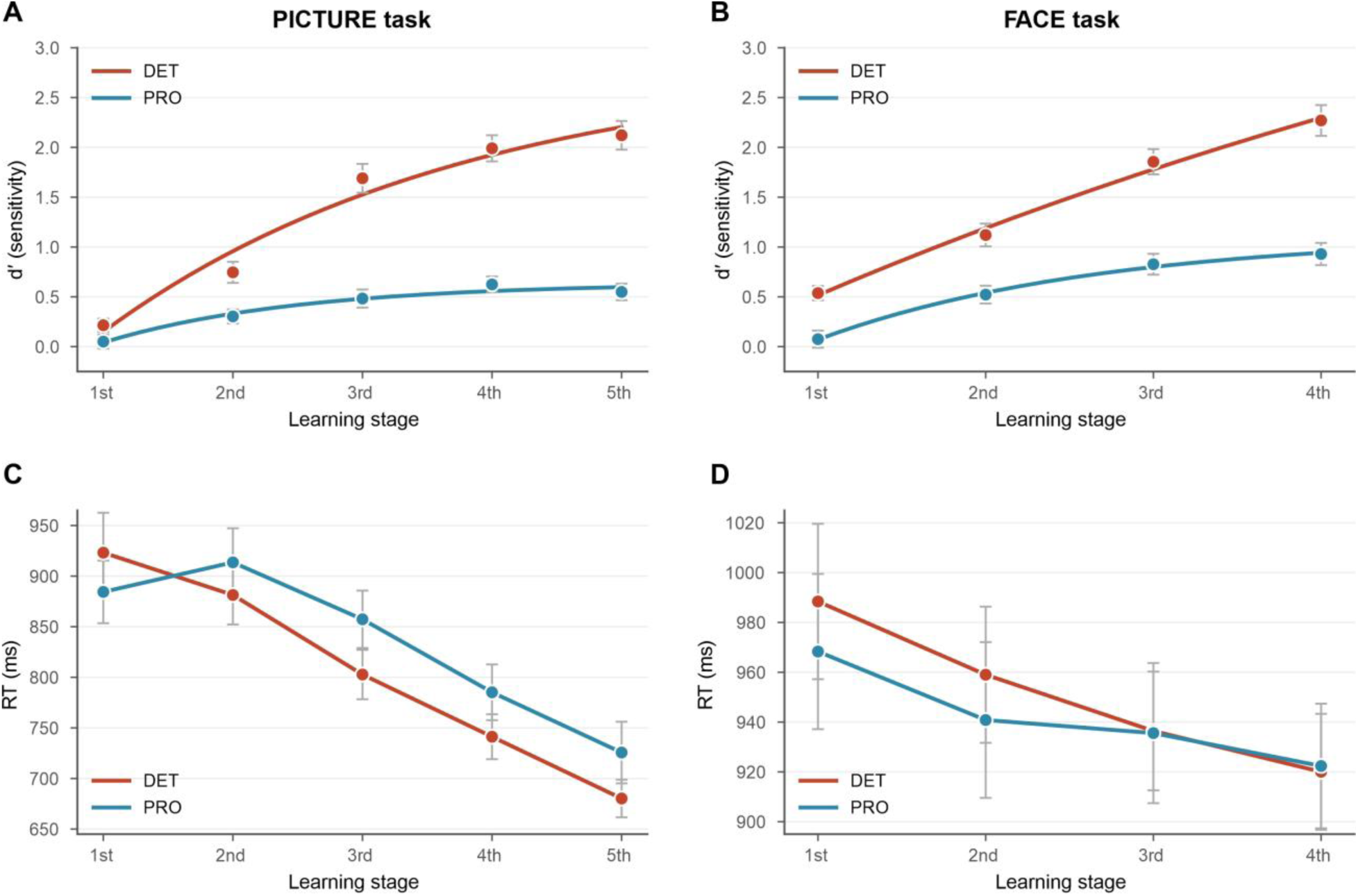
Behavioral learning performance across deterministic and probabilistic conditions. Learning accuracy (d′) (A-B) and reaction times (C-D) across learning stages in the deterministic (DET) and probabilistic (PRO) conditions for the PICTURE and FACE tasks. Error bars indicate SEM.

#### Accuracy (d′)

In the PICTURE task (Fig. 1A), there were significant main effects of Condition, F(1, 45) = 116.79, p < .001, η²ₚ = .722, and Learning Stage, F(3.29, 148.07) = 79.21, p < .001, η²ₚ = .638, as well as a significant Condition × Learning Stage interaction, F(4, 180) = 34.85, p < .001, η²ₚ = .436. Bonferroni-corrected pairwise tests confirmed that d′ in the deterministic condition exceeded that in the probabilistic condition from Stage 2 onwards (Cohen’s d increasing from 0.62 to 1.56; all p ≤ .001). In the FACE task (Fig. 1B), the same pattern was observed: significant main effects of Condition, F(1, 45) = 79.28, p < .001, η²ₚ = .638, and Learning Stage, F(2.26, 101.54) = 70.99, p < .001, η²ₚ = .612, together with a significant Condition × Learning Stage interaction, F(3, 135) = 14.67, p < .001, η²ₚ = .246. The difference between conditions was significant at every stage and increased across learning (d = 0.58–1.45).

#### Reaction time during learning

RTs decreased across learning in both tasks, although condition-dependent speeding was evident only in the PICTURE task (Fig. 1C–D). In the PICTURE task, response times shortened markedly across stages, F(1.91, 85.84) = 50.78, p < .001, η²ₚ = .530, and the Condition × Learning Stage interaction was significant, F(2.16, 97.19) = 4.76, p = .009, η²ₚ = .096, reflecting steeper speeding in the deterministic condition (923 → 680 ms) than in the probabilistic condition (884 → 726 ms). The main effect of Condition did not reach significance, F(1, 45) = 2.31, p = .136. In the FACE task, RTs decreased across stages, F(1.74, 78.32) = 6.49, p = .004, η²ₚ = .126, but neither the main effect of Condition, F(1, 45) = 0.36, p = .550, nor the Condition × Learning Stage interaction, F(2.25, 101.21) = 0.48, p = .645, was significant; both conditions reached comparable RTs by Stage 4.

In summary, participants learned the associations in both tasks, with substantially higher accuracy by the end of learning in the deterministic environment and, in the PICTURE task, faster responses. Performance also improved across learning in the probabilistic condition, indicating that the smaller behavioral gains were not simply due to a failure to engage with the task.

### Post-learning knowledge test

To verify that the associations were retained beyond performance during training, an independent post-learning test was administered separately after the DET and PRO blocks in each task. Paired t-tests on test accuracy and correct-trial RT showed better post-learning performance after deterministic than probabilistic learning in both tasks. In the FACE task, test accuracy was higher in DET than PRO (M = 0.96 vs 0.79), t(45) = 8.39, p < .001, Cohen’s d = 1.24; correct-trial RT was faster in DET than PRO (1343 vs 1548 ms), t(45) = −2.85, p = .007, d = −0.42. In the PICTURE task, test accuracy was 0.94 (DET) vs 0.71 (PRO), t(45) = 7.66, p < .001, d = 1.13; correct-trial RT was 1538 vs 1997 ms, t(45) = −5.01, p < .001, d = −0.74. Mean post-learning accuracy in the probabilistic condition exceeded the nominal chance level of 0.50 in both tasks, consistent with retention of the learned associations. At the same time, lower accuracy and slower retrieval after probabilistic than deterministic learning indicate less successful post-learning performance under partial feedback reliability. This independent post-training measure indicates that the neural differences between deterministic and probabilistic learning occurred despite substantial learning in the probabilistic condition.

Taken together, the behavioral results establish learning in both environments while showing a clear performance advantage for deterministic feedback.

### ERP analyses Response-locked: ERN

ERN was reliably larger (more negative) for error than correct trials in both tasks (Event Type main effect, both p < .001). Its learning-related trajectory, however, differed between deterministic and probabilistic conditions.

In the PICTURE task, the Condition × Event Type × Learning Stage interaction was significant, t(808) = −4.29, p < .001, qualifying lower-order Condition × Event Type and Condition × Learning Stage effects (both p < .005). As shown in Fig. 2A, ERN amplitude on error trials in the deterministic condition became progressively more negative across the five learning stages, whereas comparatively little change was observed in the probabilistic condition; ERN on correct trials remained stable in both. In the FACE task, the corresponding three-way interaction did not reach significance, t(599) = −1.88, p = .060, although the descriptive pattern was similar to the PICTURE task. The lower-order Condition × Event Type and Event Type × Learning Stage interactions were significant (p = .003 and p < .001, respectively; Fig. 2B).

**Figure 2.**
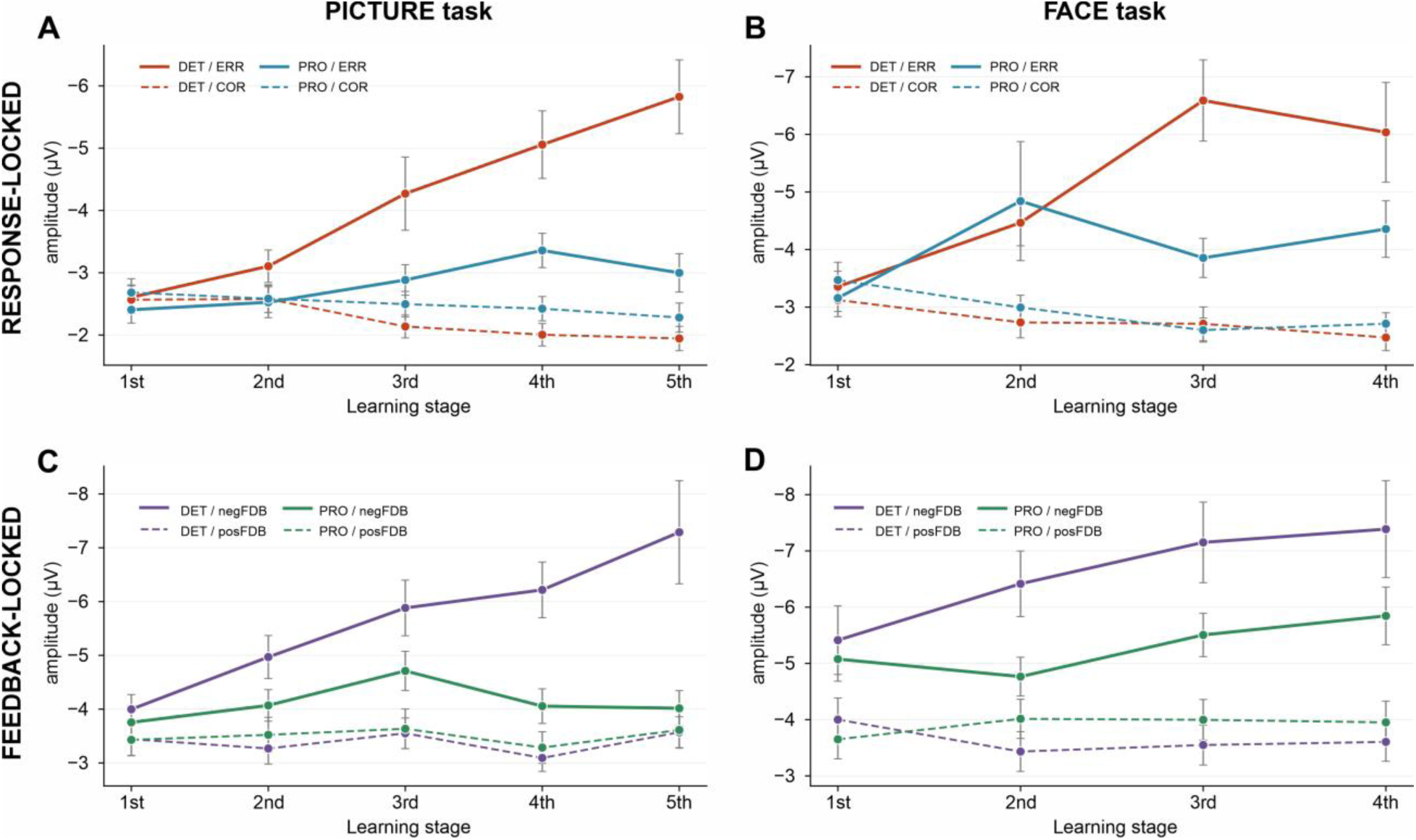
Event-related potentials dynamics across learning. Response-locked (A–B) and feedback-locked (C–D) amplitudes at FCz across learning stages in the PICTURE and FACE tasks, shown separately for deterministic (DET) and probabilistic (PRO) conditions. Solid lines indicate error/negative-feedback trials and dashed lines correct/positive-feedback trials. Error bars indicate SEM.

Thus, reliability-dependent ERN change across learning was robust in the PICTURE task, with a descriptively similar but weaker pattern in the FACE task.

### Feedback-locked: FN

The FN was reliably larger for negative than positive feedback in both tasks (Event Type main effect, both p < .001), and its learning-related trajectory also differed between conditions. In the PICTURE task, all higher-order interactions were significant, including the Condition × Event Type × Learning Stage interaction, t(808) = −3.86, p < .001 (Fig. 2C). FN amplitude on negative-feedback trials became progressively more negative in the deterministic condition while remaining comparatively stable in the probabilistic condition. In the FACE task, the Condition × Event Type interaction was robust, t(600) = −3.98, p < .001, and the Event Type × Learning Stage interaction was significant, t(600) = −2.86, p = .004; the three-way interaction did not reach significance (p = .110), although its direction was consistent with the PICTURE task (Fig. 2D).

Together, the ERP analyses showed robust reliability-dependent learning trajectories for both response- and feedback-related components in the PICTURE task, with descriptively similar but less consistently significant effects in the FACE task.

### Time–frequency analyses

Theta-band activity (4–8 Hz) was characterized using three power measures: total, phase-locked, and induced theta.

### Total theta power

Across both tasks, total theta showed fewer systematic learning-related effects than the ERPs. In the response-locked phase, neither the main effect of Condition nor the three-way interaction reached significance in either task. In the feedback-locked phase, a Condition × Learning Stage interaction was significant in both tasks (the FACE task p = .003; the PICTURE task p < .001), reflecting a decline in total theta across deterministic learning that was absent in the probabilistic condition. This pattern contrasts with the increasing ERN/FN amplitudes and motivates decomposition of total theta into its phase-locked and induced components.

### Decomposing theta: phase-locked vs induced

Total theta power was separated into phase-locked and induced components. Phase-locked theta showed a pattern similar to the ERPs: it increased across deterministic learning on error and negative-feedback trials, with comparatively little change under probabilistic learning. The Condition × Event Type interaction was significant in every task–phase combination (all p ≤ .014; Table S1). In the PICTURE task, the Condition × Event Type × Learning Stage interaction was significant in both phases (response p = .008, Fig. 3A; feedback p = .008, Fig. 3B). In the FACE task, the corresponding three-way interactions did not reach significance, although their direction was similar to the PICTURE task (Fig. 3C and Fig. 3D).

**Figure 3.**
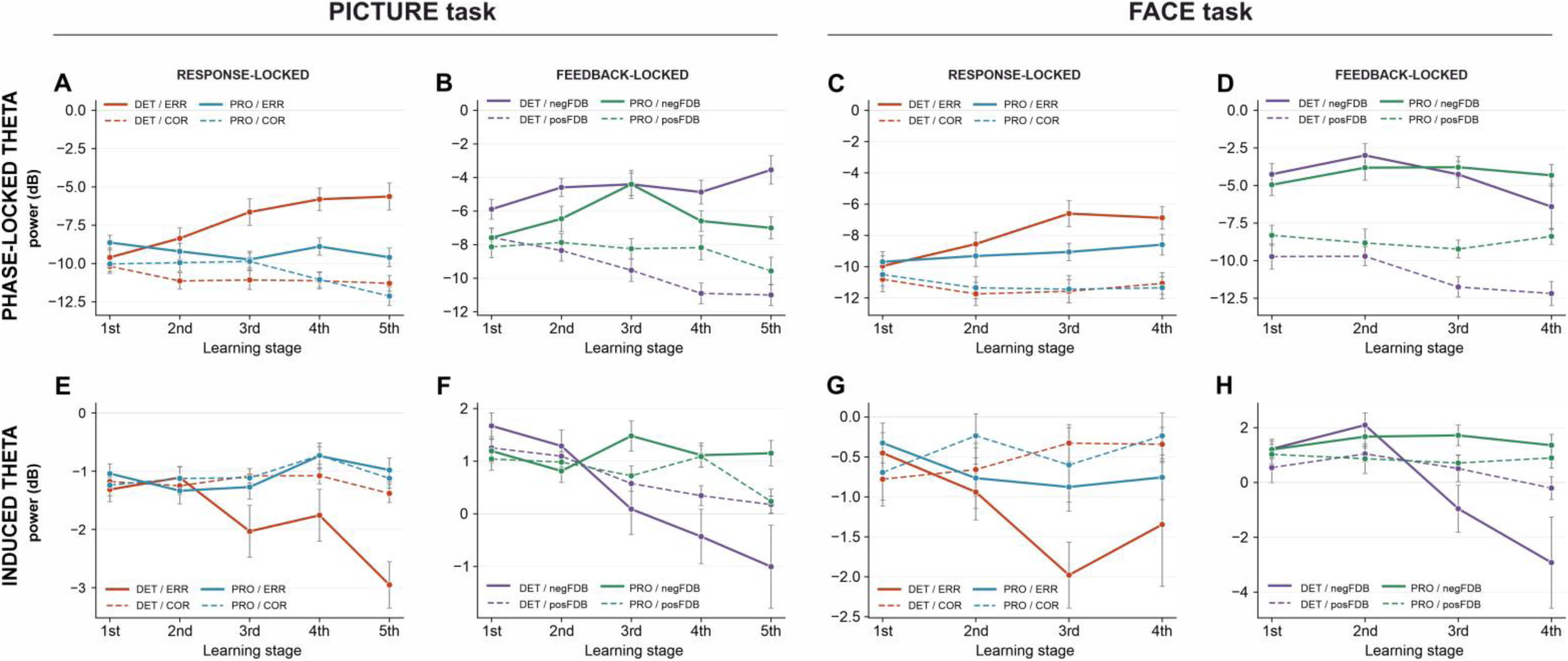
Phase-locked and induced theta dynamics across learning. Phase-locked theta power (A–D) and induced theta power (E–H) at FCz across learning stages for response-locked and feedback-locked activity in the PICTURE and FACE tasks. Solid lines indicate error/negative-feedback trials and dashed lines correct/positive-feedback trials. Power is shown in dB; error bars indicate SEM

Induced theta showed the complementary pattern, decreasing across deterministic learning on error/negative-feedback trials while remaining comparatively stable in the probabilistic condition. The Condition × Event Type × Learning Stage interaction was significant in three of four task–phase combinations (the PICTURE task response p = .004, Fig. 3E; the PICTURE task feedback p < .001, Fig. 3F; the FACE task response p = .19, Fig. 3G; the FACE task feedback p = .004, Fig. 3H).

The opposing trajectories of phase-locked and induced theta help explain the total-power results. In the response-locked phase, increases in phase-locked theta and decreases in induced theta partially offset one another, leaving total theta relatively stable. In the feedback-locked phase, the reduction in induced theta predominated over the increase in phase-locked theta, resulting in an overall decline in total theta power.

### Convergence across measures

Across measures, the clearest common pattern was observed in the PICTURE task. ERN, FN, and phase-locked theta increased across deterministic learning on error or negative-feedback trials, whereas induced theta decreased; corresponding measures remained comparatively stable under probabilistic learning. The FACE task showed broadly similar descriptive trajectories, although the Condition × Event Type × Learning Stage interaction reached significance less consistently. Total theta reflected the net contribution of the opposing phase-locked and induced dynamics: it was relatively stable in the response-locked phase, whereas in the feedback-locked phase the decrease in induced theta predominated, producing an overall decline in total power. The behavioral results provide an important context for this EEG pattern. Participants acquired and retained associations under both feedback regimes, but performance was consistently better after deterministic learning. Thus, the attenuated EEG trajectories in the probabilistic condition occurred in the presence of successful learning rather than an absence of associative acquisition.

## Discussion

The present study examined how the neural dynamics of error monitoring evolve across associative learning when the feedback signal is fully reliable as opposed to when it is partially misleading. The work builds directly on the present author’s earlier research, which established that ERN and FN deepen across deterministic learning of paired associations and that this dynamic is accompanied by progressively increasing response- and feedback-related activity in the ACC [9,10]. That earlier work was framed within the classical reinforcement-learning account of the ERN [1], in which the growth of these components reflects increasingly precise encoding of prediction errors. The present within-subject comparison between deterministic and probabilistic environments permits that interpretation to be refined.

Across the two learning paradigms, the EEG measures showed a broadly convergent pattern. All the phase-locked indices of performance monitoring (the ERN, FN and phase-locked theta) increased across deterministic learning on error and negative-feedback trials, while showing comparatively little change under probabilistic learning. These reliability-dependent dynamics were statistically robust across measures in the PICTURE task and directionally similar, although less consistently significant, in the FACE task. Induced, non-phase-locked theta showed a complementary pattern, declining across deterministic learning while remaining comparatively stable in the probabilistic condition.

The progressive deepening of ERN and FN across deterministic learning replicates our earlier findings [10] and fits the broader account of midfrontal theta as a performance-monitoring signal [2,3,8]. The novel contribution is that this dynamic depends on feedback reliability and is better characterized as a progressive tuning of the error-monitoring system than as a simple amplitude gain. When every feedback event is a valid teaching signal, complementary indices shift together: response- and feedback-locked ERPs deepen and phase-locked theta rises, while sustained non-phase-locked (induced) theta declines as performance improves. This convergence across EEG measures, two stimulus paradigms, and an independent knowledge test supports a unified account in which feedback reliability shapes how error-monitoring responses develop across learning. When 25% of feedback was misleading, this reorganization was markedly weaker or absent, even though associative learning still occurred: post-learning accuracy was lower and retrieval slower than after deterministic learning, but performance remained above chance. Midfrontal theta therefore reflects not merely the occurrence of errors but how reliably feedback can guide learning—an effect expressed through dissociable phase-locked and induced components.

This interpretation aligns with theoretical accounts of the anterior cingulate cortex as a region that integrates prediction errors with information about environmental structure [5,6], and with findings that midfrontal theta links prediction errors to behavioral adaptation [29]. Recent rodent work has provided direct causal evidence that ACC prediction-error signals contribute to adaptive updating: optogenetic silencing of ACC during task switching disrupts behavioral updating when prediction-error signals are present [30], and ACC neurons have been shown to dynamically integrate feedback information to iteratively update internal value representations [31]. The present human EEG findings extend this picture by showing that midfrontal electrophysiological signatures associated with feedback-driven updating change progressively across learning, most clearly when feedback can be trusted as a reliable learning signal. Although the present manipulation differs from outcome-volatility paradigms [32] by altering the validity of the feedback channel rather than the volatility of the underlying contingency, the conceptual implication converges: ACC-associated signals may depend not only on surprise, but also on how reliably an outcome can be used to guide subsequent behavior.

The direction of the present ERN dynamic - deepening across learning - is consistent with associative-learning paradigms, but it is worth noting that not all learning tasks produce this pattern. In motor and sequence-learning paradigms, ERN amplitude has been shown to decrease across learning as response competition diminishes with skill automatization [33]. The contrast is theoretically meaningful: in associative learning, repeated reliable feedback renders each error increasingly diagnostic of a violation against an increasingly stable expected outcome, leading to a stronger error signal; in motor learning, repeated practice reduces response conflict, leading to a weaker error signal. Both patterns reflect adaptation of the error-monitoring system to the task, but in opposing directions, and the present results align with the associative-learning literature [10,1].

The within-subject contrast also refines a purely prediction-error account of the earlier deterministic-only findings [10]. Errors and negative feedback occur in both conditions, yet the progressive tuning of phase-locked indices is most evident when feedback is fully reliable. Thus, the growth of these indices across deterministic learning appears sensitive to the reliability of feedback as a learning signal rather than to prediction-error occurrence alone. When feedback can be trusted, negative outcomes provide consistently useful information for behavioral adjustment, allowing the error-monitoring system to tune its response accordingly. This framing is compatible with work suggesting that midfrontal theta and the FN are sensitive to the salience of feedback events as well as to their value [17,18]. The earlier prediction-error interpretation therefore remains relevant, but the present data place it within a broader account in which feedback reliability modulates how strongly error-monitoring responses are tuned over learning. Although deterministic versus probabilistic learning has been examined previously using feedback-validity manipulations [11,12,13,14], those studies typically used the contrast as a secondary axis for probing individual differences, particularly aging, with learning dynamics often reduced to an early-versus-late comparison. The present design treats the trajectory across multiple within-subject learning stages as the central phenomenon and feedback reliability as the within-subject modulator of that trajectory. This design reveals that the deterministic– probabilistic difference is not merely a static condition effect, but develops across learning, becoming particularly evident at later stages when deterministic feedback has repeatedly confirmed the underlying associations while probabilistic feedback remains partially uninformative.

A central methodological contribution of the present study is the dissociation between phase-locked and induced theta sub-components, which suggests that the deterministic learning trajectory reflects complementary changes rather than a single neural process. In feedback-locked epochs, total theta power declined across deterministic learning even as FN amplitude deepened, a pattern that is difficult to interpret from total theta alone. Decomposing theta into a phase-locked component, which contributes to the trial-averaged ERP, and an induced component, whose phase varies across trials, reveals two distinct trajectories. Phase-locked theta increased across deterministic learning on error and negative-feedback trials and remained comparatively stable in the probabilistic condition, whereas induced theta declined across deterministic learning and remained comparatively stable under probabilistic learning. This theta decomposition also clarifies the differing total-theta trajectories across epoch types. In response-locked epochs, opposing changes in phase-locked and induced theta partially offset one another, leaving total theta relatively stable. In feedback-locked epochs, by contrast, the decline in induced theta outweighed the increase in phase-locked theta, producing an overall reduction in total theta power. Thus, total theta alone can mask distinct and opposing sub-component dynamics.

The convergence of ERPs (ERN, FN) with phase-locked theta is grounded in the foundational observation that the ERN is, to a large extent, a time-domain expression of phase-locked midfrontal theta, with the two measures reflecting closely related neural activity at different temporal resolutions [15,16]. Phase-locked / non-phase-locked decomposition has been further developed methodologically in feedback-processing research using time-frequency analyses of gambling and reward paradigms [21,19]. Particularly relevant is work in extensive motor practice showing that induced (non-phase-locked) frontal theta decreases across training while evoked theta and the FN do not show the same reduction [20]. The present results extend this dissociation to associative learning and suggest that its expression depends on the reliability of feedback as a teaching signal.

The two sub-components may reflect complementary aspects of learning-related tuning under deterministic feedback. The progressive increase in phase-locked theta, together with the increasing ERN and FN amplitudes, is consistent with stronger event-locked responses to errors and negative feedback as deterministic learning progresses. The concurrent decline in induced theta may reflect reduced sustained, non-phase-locked engagement as the task becomes more familiar and less demanding. These changes need not reflect opposing processes: greater event-specific differentiation can coexist with reduced generalized engagement [34,17].

The overall direction of effects was similar across two associative-learning paradigms with markedly different stimulus materials—image–image pairs and face–name pairs—providing evidence for cross-paradigm consistency rather than a stimulus-specific pattern. The inferential evidence was nevertheless substantially stronger in the PICTURE task, where the central Condition × Response × Stage interaction was significant for ERN, FN, phase-locked theta, and induced theta, whereas in the FACE task several corresponding three-way interactions did not reach significance despite generally similar directions of effect. The smaller number of learning stages in the FACE task (four, compared to five in the PICTURE task) may have reduced sensitivity to longitudinal interactions, although other differences between the tasks may also have contributed. Accordingly, the present findings support a common qualitative pattern across paradigms, but they should not be interpreted as demonstrating statistical equivalence of the effects across tasks.

Several limitations should be noted. First, the present sample consisted of healthy younger adults, and the generalizability of these dynamics to clinical, developmental, or older populations remains to be established. Second, the contrast between 100% and 75% feedback reliability operationalizes feedback validity at only two fixed levels; a parametric manipulation across a wider range of validity levels would more directly test whether the neural sensitivity to reliability is continuous or categorical. Finally, because the present analyses treat the two tasks separately, future work could test cross-task generalization more directly in a joint model.

The reliability dependence of error-monitoring dynamics observed here raises testable hypotheses for populations in which feedback-driven learning is impaired. Older adults are known to show greater difficulty in probabilistic than deterministic learning [11,35]; the present results suggest that future work could examine whether this behavioural difference is accompanied by altered development of the phase-locked learning trajectory observed here. More broadly, the phase-locked / induced dissociation offers a finer-grained framework for asking whether atypical feedback processing is associated primarily with altered event-locked error responses, sustained cognitive engagement, or both.

The present findings also raise the possibility that feedback reliability may be particularly important during early acquisition, a hypothesis that could be tested directly in educational and rehabilitation settings rather than assumed from the current design. More broadly, the data suggests that the error-monitoring system is sensitive not simply to individual feedback events but to the reliability of the environment from which those events arise. This perspective recasts midfrontal theta from a generic error signal toward a context-sensitive index of how consistently feedback can be used as a basis for learning.

## Methods

### Participants

Forty-six adults (25 male, 21 female; age range 19–26 years, M = 21.7, SD = 2.1) participated in the study. Participants were recruited via university notice boards and online advertisements (Facebook, posters at the Jagiellonian University, Kraków) and received monetary compensation for their participation. All participants were right-handed, had normal or corrected-to-normal vision, and were free from any current or past psychiatric or neurological conditions, developmental disorders (including attention-deficit/hyperactivity disorder and dyslexia), and current use of psychoactive substances. All participants provided written informed consent prior to the experiment. Three participants were excluded from all EEG analyses due to systematic technical artifacts (extreme amplitude values indicative of channel or reference-related problems). The study was approved by the Research Ethics Committee of the Faculty of Management and Social Communication, Jagiellonian University and was conducted in accordance with the Declaration of Helsinki.

### General study structure

Each participant completed two associative-learning tasks during a single EEG session: a paired-associate image–image task (the PICTURE task), adapted from Gawlowska et al. [9], and a face– name task (the FACE task), adapted from Alm et al. [23]. Both tasks were administered in two predictability conditions: a deterministic condition (DET) in which feedback was fully reliable (100% predictive), and a probabilistic condition (PRO) in which feedback was reliable on 75% of trials. Participants were informed at the outset of the probabilistic blocks that 25% of the feedback in those blocks would be misleading. The predictability of the upcoming block was further signaled visually by a colored frame surrounding each trial: a violet frame for the deterministic blocks and a yellow frame for the probabilistic blocks.

The order of the two tasks (PICTURE first vs. FACE first) was counterbalanced across participants. The order of the two predictability conditions (DET first vs. PRO first) within each task was also independently counterbalanced. This double counterbalancing controlled for fatigue, practice, and order effects at both levels (between-task and between-condition).

### Image–image associative learning task (the PICTURE task)

The PICTURE task was adapted from Gawlowska et al. [9] and used the same set of stimuli - everyday-object photographs on white backgrounds [36]. The task required participants to learn fixed associations between pairs of images. Within each predictability condition, nine target pairs (e.g. remote control — raspberry) were designated as correct. For each target pair, three foil pairs were constructed by recombining elements across triads — for example, remote control — strawberry, raspberry — cassette, and cassette — strawberry. This yielded 36 distinct trial types per condition (9 correct + 27 foil pairs), of which one quarter were correct and three quarters were foils.

Each learning block consisted of five learning stages. Within each stage, all 36 trial types were presented once in randomized order; participants thus encountered each correct pair five times and each foil pair five times across the block (180 trials in total per condition). On each trial, the pair of images was first displayed for 1,500 ms (with the colored frame indicating the feedback-reliability condition), followed by a 1,000-ms response cue during which the images remained visible together with placeholders for the two response options (YES/NO); participants were instructed to indicate whether the pair was correct or not. Responses were collected until 2,500 ms after the onset of the response cue. After the response (or after the response window had elapsed), a blank screen appeared for approximately 1,000 ms and then feedback was presented for approximately 1,000 ms, indicating to the participant whether the response had been correct (GOOD) or not (BAD). Feedback was followed by a blank screen of approximately 1,000 ms. In the deterministic condition the feedback was always veridical; in the probabilistic condition the feedback was veridical on 75% of trials and misleading on 25% of trials.

The post-learning knowledge test, administered immediately after the training block, used a two-alternative forced-choice (2AFC) recognition format that differed from the training. On each test trial, a single target image (the cue) was displayed together with two response-option images arranged vertically and labelled “1” and “2”; one option was the correct paired associate from the training set and the other was a distractor drawn from a foil pair. The cue appeared on either the left or right side of the screen with the response options on the opposite side, with side counterbalanced across trials. Participants pressed the corresponding key to indicate which of the two options had been paired with the cue during training. No feedback was provided during the test. Each of the nine target pairs was probed twice, once with each element of the pair serving as the cue, yielding 18 test trials per condition.

### Face–name associative learning task (FACE task)

The FACE task was adapted from Alm et al. [23] and required participants to learn fixed associations between photographs of unfamiliar faces and Polish first names. Face stimuli were obtained from the Face Research Lab London Set [37] and depicted eight faces with neutral expression, balanced for gender (four female and four male); the associated names were common Polish first names matched to the apparent gender of each face. Within each predictability condition, eight target face–name pairs were designated as correct. For each target face, three foil names were used — different from the target’s correct name. This yielded 32 distinct trial types per condition (8 correct + 24 foil pairings).

Each learning block consisted of four learning stages. Within each stage, all 32 trial types were presented once in randomized order; participants thus encountered each correct pairing four times and each foil pairing four times across the block (128 trials in total per condition). On each trial, the face with the candidate name printed below it was first displayed for 1,500 ms (with the colored frame indicating the feedback-reliability condition), followed by a 500-ms response cue during which the face–name display remained visible with placeholders for the two response options (YES/NO); participants were instructed to indicate whether the name matched the face according to their learning history. Responses were collected until 2,500 ms after the onset of the response cue. After the response (or after the response window had elapsed), a blank screen appeared for approximately 1,000 ms and then feedback was presented for approximately 1,000 ms, indicating to the participant whether the response had been correct (GOOD) or not (BAD). Feedback was followed by a blank screen of approximately 1,000 ms. As in the PICTURE task, feedback was always veridical in the deterministic condition and was veridical on 75% of trials in the probabilistic condition.

The post-learning knowledge test, administered immediately after the training block, used the same two-alternative forced-choice format as in the PICTURE task. On each test trial, a face served as the cue and two candidate names appeared as response options labelled “1” and “2”; one name was the correct learned associate, and the other was a name from a different target face. The cue–options spatial layout and side counterbalancing matched the PICTURE task test. No feedback was provided. Each of the eight faces was probed twice with different distractor names, yielding 16 test trials per condition.

### EEG recording

Continuous dense-array EEG was recorded from 256 channels using a HydroCel Geodesic Sensor Net (EGI System 300; Electrical Geodesic Inc., OR, USA) at a sampling rate of 1000 Hz, with an online band-pass filter of 0.01–100 Hz and the vertex electrode as the reference. Data was recorded using NetStation Software (Version 4.5.1, Electrical Geodesic Inc., OR, USA). Electrode impedances were kept below 50 kΩ. Offline analyses were conducted using the open-source EEGLAB toolbox [38].

### EEG preprocessing

Channels located on the face were removed before any preprocessing, leaving 224 scalp channels for all subsequent processing and analyses. The continuous data was band-pass filtered between 0.5 and 35 Hz (zero-phase FIR filter, applied separately as a 0.5-Hz high-pass and a 35-Hz low-pass). Bad channels were automatically removed by kurtosis measures with a threshold value of 5 standard deviations (on average 10.3% of the channels). Continuous data was then visually inspected in order to manually remove remaining bad channels or time epochs containing high-amplitude, high-frequency muscle noise and other irregular artifacts. Average reference was recomputed. Independent component analysis was used to remove artifacts from the data. Due to the large number of channels, decomposition of EEG data with the Infomax algorithm was preceded with Principal Component Analysis. Fifty independent components were extracted and visually inspected for each subject. On the basis of the spatiotemporal pattern, components recognized as blinks, heart rate, saccades, muscle artifacts, or bad channels were removed [39,40]. Missing channels were reconstructed using spherical interpolation.

### EEG analysis

#### Event-related potentials (ERPs)

For the ERP analyses, the EEG data was segmented separately around response onset and feedback onset. Response-locked epochs extended from −500 to +1000 ms relative to the response and were baseline-corrected using the −300 to −100 ms pre-response interval. Feedback-locked epochs extended from −500 to +1500 ms relative to feedback onset and were baseline-corrected using the −200 to 0 ms pre-feedback interval.

Following previously applied methodology [41,42,10,43], post-response and post-feedback amplitudes were quantified using peak-to-peak measures. The ERN was quantified at the FCz electrode as the amplitude difference between the most negative peak occurring up to 100 ms after an incorrect response and the most positive peak within the 100 ms interval before the response onset. The same time windows were used for epochs corresponding to correct responses. The FN was quantified at the FCz electrode as the amplitude difference between the most negative peak occurring 200–350 ms after negative feedback and the most positive peak within the 100–200 ms post-feedback interval. The same time windows were applied to epochs containing positive feedback.

#### Time–frequency analyses

Time–frequency analyses were performed on the same response-locked and feedback-locked epochs using complex Morlet wavelets. The decomposition included 20 logarithmically spaced frequencies from 4 to 12 Hz, with a wavelet width of four cycles. From the resulting wavelet coefficients, three theta-band (4–8 Hz) power measures were derived [19]: total theta power, phase-locked theta power, and induced theta power.

Total theta power was calculated as the trial-averaged squared magnitude of the wavelet coefficients. Phase-locked (evoked) theta power was calculated by averaging the complex wavelet coefficients across trials and subsequently squaring their magnitude, thereby capturing activity that was phase-consistent across trials. Induced (non-phase-locked) theta power was calculated as the difference between total and phase-locked power in linear units prior to conversion to decibels. All three power measures were converted to decibels (dB) relative to a common −500 to 0 ms baseline and subsequently averaged across the theta-frequency range. The same baseline was used for all three measures to ensure comparability while preserving the relationship between total, phase-locked, and induced power in linear units.

For each measure, values were averaged within predefined time windows based on previous literature on midfrontal theta activity during performance monitoring [2]. Response-locked measures were averaged over the 0–300 ms post-response interval, whereas feedback-locked measures were averaged over the 200–400 ms post-feedback interval.

### Statistical analysis

Behavioral measures, including d′ and reaction times during learning, were analyzed using repeated-measures ANOVAs for the within-subject factors of Condition and Learning Stage. Sphericity was assessed using Mauchly’s test. When the assumption of sphericity was violated, Greenhouse–Geisser-corrected degrees of freedom and p values were reported. Post hoc pairwise comparisons were adjusted using the Bonferroni correction. Post-learning test performance and reaction times were compared between conditions using paired-samples t-tests. The d′ measure was calculated according to signal detection theory [27], with the Hautus log-linear correction [28] applied in cases of extreme hit or false-alarm rates.

EEG measures were analyzed using linear mixed-effects models. Separate models were fitted for each EEG measure (ERN, FN, total theta, phase-locked theta, and induced theta) within each task. Each model included Condition (DET vs. PRO), Event Type (error vs. correct responses for response-locked measures; negative vs. positive feedback for feedback-locked measures), Learning Stage, and all interactions among these predictors, with a random intercept for participant:

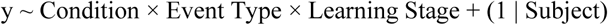

Models were fitted using restricted maximum likelihood (REML). Sum coding was used for the categorical predictors to facilitate interpretation of main effects in the presence of interactions. Deterministic and probabilistic conditions were coded as +0.5 and −0.5, respectively; error responses or negative feedback were coded as +0.5, whereas correct responses or positive feedback were coded as −0.5.

Learning Stage was entered as a continuous, mean-centered linear predictor rather than as a categorical factor. Consequently, its main effect and interactions with Condition and Event Type represent the linear rate of change across learning stages. This parameterization directly addresses whether each EEG measure changes progressively over the course of learning. Model coefficients are reported as β with standard errors (SE), together with t statistics, degrees of freedom, and p values.

All statistical analyses were performed using IBM SPSS Statistics (Version 29; IBM Corp., 2022) and MATLAB (R2019b; The MathWorks, Inc., 2019).

### Use of generative AI

Generative artificial intelligence (ChatGPT, OpenAI) was used during manuscript preparation to assist with language editing, clarity, consistency, manuscript organization, and formatting. The author critically reviewed and revised all AI-assisted text and takes full responsibility for the content of the manuscript.

## Acknowledgements

The author gratefully acknowledges Dominika Fortuna, MA, for her valuable assistance with participant recruitment and data collection.

## Author contributions

M.G. designed the study, acquired and analyzed the data, interpreted the results, and wrote and revised the manuscript.

## Data Availability

The datasets generated and analyzed during the current study are available from the corresponding author upon reasonable request.

