## Supplementary Table 1 for "Feedback reliability modulates learning-related dynamics of midfrontal error monitoring"

Supplementary Table S1. Fixed-effect estimates from linear mixed-effects models of EEG measures.

| **Measure** | **Task** | **locking** | **Effect** | **β** | **SE** | **t** | **df** | **p** |
|---|---|---|---|---|---|---|---|---|
| ERN | FACE | response-locked | Condition | -0,444 | 0,214 | -2,07 | 599 | 0.039 |
| ERN | FACE | response-locked | Event Type | -1,731 | 0,214 | -8,07 | 599 | <.001 |
| ERN | FACE | response-locked | Learning Stage | -0,214 | 0,096 | -2,22 | 599 | 0.027 |
| ERN | FACE | response-locked | Condition × Event Type | -1,258 | 0,429 | -2,93 | 599 | 0.003 |
| ERN | FACE | response-locked | Condition × Learning Stage | -0,431 | 0,193 | -2,24 | 599 | 0.025 |
| ERN | FACE | response-locked | Event Type × Learning Stage | -0,893 | 0,193 | -4,63 | 599 | <.001 |
| ERN | FACE | response-locked | Condition × Event Type × Learning Stage | -0,725 | 0,385 | -1,88 | 599 | 0.060 |
| ERN | PICTURE | response-locked | Condition | -0,549 | 0,119 | -4,61 | 808 | <.001 |
| ERN | PICTURE | response-locked | Event Type | -1,137 | 0,119 | -9,54 | 808 | <.001 |
| ERN | PICTURE | response-locked | Learning Stage | -0,191 | 0,042 | -4,53 | 808 | <.001 |
| ERN | PICTURE | response-locked | Condition × Event Type | -1,591 | 0,238 | -6,68 | 808 | <.001 |
| ERN | PICTURE | response-locked | Condition × Learning Stage | -0,276 | 0,084 | -3,28 | 808 | 0.001 |
| ERN | PICTURE | response-locked | Event Type × Learning Stage | -0,66 | 0,084 | -7,83 | 808 | <.001 |
| ERN | PICTURE | response-locked | Condition × Event Type × Learning Stage | -0,723 | 0,168 | -4,29 | 808 | <.001 |
| FN | FACE | feedback-locked | Condition | -0,509 | 0,192 | -2,66 | 600 | 0.008 |
| FN | FACE | feedback-locked | Event Type | -2,154 | 0,192 | -11,24 | 600 | <.001 |
| FN | FACE | feedback-locked | Learning Stage | -0,236 | 0,086 | -2,75 | 600 | 0.006 |
| FN | FACE | feedback-locked | Condition × Event Type | -1,525 | 0,383 | -3,98 | 600 | <.001 |
| FN | FACE | feedback-locked | Condition × Learning Stage | -0,079 | 0,172 | -0,46 | 600 | 0.644 |
| FN | FACE | feedback-locked | Event Type × Learning Stage | -0,491 | 0,172 | -2,86 | 600 | 0.004 |
| FN | FACE | feedback-locked | Condition × Event Type × Learning Stage | -0,551 | 0,344 | -1,6 | 600 | 0.110 |
| FN | PICTURE | feedback-locked | Condition | -0,723 | 0,135 | -5,35 | 808 | <.001 |
| FN | PICTURE | feedback-locked | Event Type | -1,459 | 0,135 | -10,8 | 808 | <.001 |
| FN | PICTURE | feedback-locked | Learning Stage | -0,214 | 0,048 | -4,48 | 808 | <.001 |
| FN | PICTURE | feedback-locked | Condition × Event Type | -1,67 | 0,27 | -6,18 | 808 | <.001 |
| FN | PICTURE | feedback-locked | Condition × Learning Stage | -0,364 | 0,095 | -3,81 | 808 | <.001 |
| FN | PICTURE | feedback-locked | Event Type × Learning Stage | -0,406 | 0,095 | -4,26 | 808 | <.001 |
| FN | PICTURE | feedback-locked | Condition × Event Type × Learning Stage | -0,737 | 0,191 | -3,86 | 808 | <.001 |
| Total theta | FACE | feedback-locked | Condition | -0,593 | 0,248 | -2,4 | 617 | 0.017 |

| | | | | | | | | |
|---|---|---|---|---|---|---|---|---|
| Total theta | FACE | feedback-locked | Event Type | 1,231 | 0,248 | 4,97 | 617 | <.001 |
| Total theta | FACE | feedback-locked | Learning Stage | -0,267 | 0,112 | -2,39 | 617 | 0.017 |
| Total theta | FACE | feedback-locked | Condition × Event Type | -0,404 | 0,495 | -0,82 | 617 | 0.415 |
| Total theta | FACE | feedback-locked | Condition × Learning Stage | -0,661 | 0,223 | -2,96 | 617 | 0.003 |
| Total theta | FACE | feedback-locked | Event Type × Learning Stage | -0,059 | 0,223 | -0,26 | 617 | 0.792 |
| Total theta | FACE | feedback-locked | Condition × Event Type × Learning Stage | -0,872 | 0,447 | -1,95 | 617 | 0.051 |
| Total theta | FACE | response-locked | Condition | 0,068 | 0,123 | 0,55 | 616 | 0.581 |
| Total theta | FACE | response-locked | Event Type | 0,174 | 0,123 | 1,41 | 616 | 0.158 |
| Total theta | FACE | response-locked | Learning Stage | 0,036 | 0,056 | 0,65 | 616 | 0.518 |
| Total theta | FACE | response-locked | Condition × Event Type | 0,015 | 0,246 | 0,06 | 616 | 0.951 |
| Total theta | FACE | response-locked | Condition × Learning Stage | 0,01 | 0,111 | 0,09 | 616 | 0.932 |
| Total theta | FACE | response-locked | Event Type × Learning Stage | -0,031 | 0,111 | -0,28 | 616 | 0.781 |
| Total theta | FACE | response-locked | Condition × Event Type × Learning Stage | 0,341 | 0,222 | 1,53 | 616 | 0.126 |
| Total theta | PICTURE | feedback-locked | Condition | -0,192 | 0,118 | -1,63 | 788 | 0.103 |
| Total theta | PICTURE | feedback-locked | Event Type | 0,652 | 0,118 | 5,55 | 788 | <.001 |
| Total theta | PICTURE | feedback-locked | Learning Stage | -0,21 | 0,042 | -5,06 | 788 | <.001 |
| Total theta | PICTURE | feedback-locked | Condition × Event Type | 0,105 | 0,235 | 0,44 | 788 | 0.657 |
| Total theta | PICTURE | feedback-locked | Condition × Learning Stage | -0,313 | 0,083 | -3,76 | 788 | <.001 |
| Total theta | PICTURE | feedback-locked | Event Type × Learning Stage | 0,087 | 0,083 | 1,05 | 788 | 0.295 |
| Total theta | PICTURE | feedback-locked | Condition × Event Type × Learning Stage | -0,187 | 0,166 | -1,13 | 788 | 0.261 |
| Total theta | PICTURE | response-locked | Condition | -0,029 | 0,095 | -0,3 | 788 | 0.763 |
| Total theta | PICTURE | response-locked | Event Type | 0,37 | 0,095 | 3,91 | 788 | <.001 |
| Total theta | PICTURE | response-locked | Learning Stage | 0,008 | 0,033 | 0,25 | 788 | 0.806 |
| Total theta | PICTURE | response-locked | Condition × Event Type | 0,326 | 0,189 | 1,72 | 788 | 0.085 |
| Total theta | PICTURE | response-locked | Condition × Learning Stage | -0,02 | 0,067 | -0,3 | 788 | 0.762 |
| Total theta | PICTURE | response-locked | Event Type × Learning Stage | 0,082 | 0,067 | 1,23 | 788 | 0.219 |
| Total theta | PICTURE | response-locked | Condition × Event Type × Learning Stage | 0,046 | 0,134 | 0,34 | 788 | 0.732 |
| Phase-locked theta | FACE | feedback-locked | Condition | -1,123 | 0,323 | -3,48 | 617 | <.001 |
| Phase-locked theta | FACE | feedback-locked | Event Type | 5,461 | 0,323 | 16,92 | 617 | <.001 |
| Phase-locked theta | FACE | feedback-locked | Learning Stage | -0,353 | 0,146 | -2,42 | 617 | 0.016 |
| Phase-locked theta | FACE | feedback-locked | Condition × Event Type | 2,005 | 0,646 | 3,11 | 617 | 0.002 |
| Phase-locked theta | FACE | feedback-locked | Condition × Learning Stage | -0,835 | 0,291 | -2,87 | 617 | 0.004 |
| Phase-locked theta | FACE | feedback-locked | Event Type × Learning Stage | 0,298 | 0,291 | 1,02 | 617 | 0.307 |
| Phase-locked theta | FACE | feedback-locked | Condition × Event Type × Learning Stage | 0,087 | 0,583 | 0,15 | 617 | 0.881 |

| | | | | | | | | |
|---|---|---|---|---|---|---|---|---|
| Phase-locked theta | FACE | response-locked | Condition | 0,558 | 0,286 | 1,95 | 616 | 0.052 |
| Phase-locked theta | FACE | response-locked | Event Type | 2,674 | 0,286 | 9,34 | 616 | <.001 |
| Phase-locked theta | FACE | response-locked | Learning Stage | 0,321 | 0,129 | 2,48 | 616 | 0.013 |
| Phase-locked theta | FACE | response-locked | Condition × Event Type | 1,407 | 0,572 | 2,46 | 616 | 0.014 |
| Phase-locked theta | FACE | response-locked | Condition × Learning Stage | 0,553 | 0,258 | 2,14 | 616 | 0.033 |
| Phase-locked theta | FACE | response-locked | Event Type × Learning Stage | 0,966 | 0,258 | 3,74 | 616 | <.001 |
| Phase-locked theta | FACE | response-locked | Condition × Event Type × Learning Stage | 0,704 | 0,517 | 1,36 | 616 | 0.174 |
| Phase-locked theta | PICTURE | feedback-locked | Condition | 0,337 | 0,256 | 1,32 | 788 | 0.188 |
| Phase-locked theta | PICTURE | feedback-locked | Event Type | 3,394 | 0,256 | 13,27 | 788 | <.001 |
| Phase-locked theta | PICTURE | feedback-locked | Learning Stage | -0,177 | 0,09 | -1,96 | 788 | 0.051 |
| Phase-locked theta | PICTURE | feedback-locked | Condition × Event Type | 2,814 | 0,512 | 5,5 | 788 | <.001 |
| Phase-locked theta | PICTURE | feedback-locked | Condition × Learning Stage | -0,14 | 0,181 | -0,78 | 788 | 0.438 |
| Phase-locked theta | PICTURE | feedback-locked | Event Type × Learning Stage | 0,905 | 0,181 | 5 | 788 | <.001 |
| Phase-locked theta | PICTURE | feedback-locked | Condition × Event Type × Learning Stage | 0,965 | 0,362 | 2,67 | 788 | 0.008 |
| Phase-locked theta | PICTURE | response-locked | Condition | 0,822 | 0,238 | 3,45 | 788 | <.001 |
| Phase-locked theta | PICTURE | response-locked | Event Type | 2,571 | 0,238 | 10,79 | 788 | <.001 |
| Phase-locked theta | PICTURE | response-locked | Learning Stage | 0,035 | 0,084 | 0,42 | 788 | 0.677 |
| Phase-locked theta | PICTURE | response-locked | Condition × Event Type | 2,37 | 0,477 | 4,97 | 788 | <.001 |
| Phase-locked theta | PICTURE | response-locked | Condition × Learning Stage | 0,76 | 0,168 | 4,51 | 788 | <.001 |
| Phase-locked theta | PICTURE | response-locked | Event Type × Learning Stage | 0,821 | 0,168 | 4,87 | 788 | <.001 |
| Phase-locked theta | PICTURE | response-locked | Condition × Event Type × Learning Stage | 0,903 | 0,337 | 2,68 | 788 | 0.008 |
| Induced theta | FACE | feedback-locked | Condition | -0,934 | 0,241 | -3,88 | 617 | <.001 |
| Induced theta | FACE | feedback-locked | Event Type | 0,07 | 0,241 | 0,29 | 617 | 0.773 |
| Induced theta | FACE | feedback-locked | Learning Stage | -0,426 | 0,109 | -3,92 | 617 | <.001 |
| Induced theta | FACE | feedback-locked | Condition × Event Type | -1,079 | 0,481 | -2,24 | 617 | 0.025 |
| Induced theta | FACE | feedback-locked | Condition × Learning Stage | -0,848 | 0,217 | -3,91 | 617 | <.001 |
| Induced theta | FACE | feedback-locked | Event Type × Learning Stage | -0,52 | 0,217 | -2,39 | 617 | 0.017 |
| Induced theta | FACE | feedback-locked | Condition × Event Type × Learning Stage | -1,252 | 0,434 | -2,88 | 617 | 0.004 |
| Induced theta | FACE | response-locked | Condition | -0,309 | 0,151 | -2,05 | 616 | 0.041 |
| Induced theta | FACE | response-locked | Event Type | -0,454 | 0,151 | -3 | 616 | 0.003 |
| Induced theta | FACE | response-locked | Learning Stage | -0,08 | 0,068 | -1,17 | 616 | 0.242 |
| Induced theta | FACE | response-locked | Condition × Event Type | -0,444 | 0,302 | -1,47 | 616 | 0.143 |
| Induced theta | FACE | response-locked | Condition × Learning Stage | -0,118 | 0,136 | -0,86 | 616 | 0.388 |
| Induced theta | FACE | response-locked | Event Type × Learning Stage | -0,425 | 0,137 | -3,11 | 616 | 0.002 |

| | | | | | | | | |
|---|---|---|---|---|---|---|---|---|
| Induced theta | FACE | response-locked | Condition × Event Type × Learning Stage | -0,362 | 0,273 | -1,33 | 616 | 0.185 |
| Induced theta | PICTURE | feedback-locked | Condition | -0,467 | 0,114 | -4,08 | 770 | <.001 |
| Induced theta | PICTURE | feedback-locked | Event Type | -0,015 | 0,114 | -0,13 | 770 | 0.893 |
| Induced theta | PICTURE | feedback-locked | Learning Stage | -0,283 | 0,041 | -6,97 | 770 | <.001 |
| Induced theta | PICTURE | feedback-locked | Condition × Event Type | -0,693 | 0,229 | -3,03 | 770 | 0.003 |
| Induced theta | PICTURE | feedback-locked | Condition × Learning Stage | -0,436 | 0,081 | -5,37 | 770 | <.001 |
| Induced theta | PICTURE | feedback-locked | Event Type × Learning Stage | -0,124 | 0,081 | -1,53 | 770 | 0.127 |
| Induced theta | PICTURE | feedback-locked | Condition × Event Type × Learning Stage | -0,592 | 0,162 | -3,65 | 770 | <.001 |
| Induced theta | PICTURE | response-locked | Condition | -0,438 | 0,092 | -4,78 | 770 | <.001 |
| Induced theta | PICTURE | response-locked | Event Type | -0,32 | 0,092 | -3,49 | 770 | <.001 |
| Induced theta | PICTURE | response-locked | Learning Stage | -0,069 | 0,033 | -2,14 | 770 | 0.033 |
| Induced theta | PICTURE | response-locked | Condition × Event Type | -0,625 | 0,183 | -3,41 | 770 | <.001 |
| Induced theta | PICTURE | response-locked | Condition × Learning Stage | -0,274 | 0,065 | -4,21 | 770 | <.001 |
| Induced theta | PICTURE | response-locked | Event Type × Learning Stage | -0,178 | 0,065 | -2,74 | 770 | 0.006 |
| Induced theta | PICTURE | response-locked | Condition × Event Type × Learning Stage | -0,375 | 0,13 | -2,88 | 770 | 0.004 |
